# Quantification of horizontal and vertical distribution of junctional proteins in fixed epithelial cells

**DOI:** 10.1101/2025.05.01.651687

**Authors:** Madeline Lovejoy, Andoni Mucci, Agustín Rabino, Rafael Garcia-Mata

## Abstract

Polarized epithelial cells form a tightly packed monolayer where individual cells are connected by cell-cell junctions, including tight junctions (TJ) and adherens junctions (AJ). Here, we present techniques for quantifying the horizontal and vertical distribution of junctional proteins in confluent, fixed epithelial cells. This approach is utilized to evaluate variations in the intensity and localization of the proteins that compose the AJ and TJ under different experimental conditions. Although our protocol is optimized for Madin-Darby Canine Kidney (MDCK) cells, it is adaptable to any cell line capable of forming cell-cell junctions.

For complete details on the use and execution of this protocol, please refer to Rabino et al. (2024). ^1^

**Graphical abstract:** 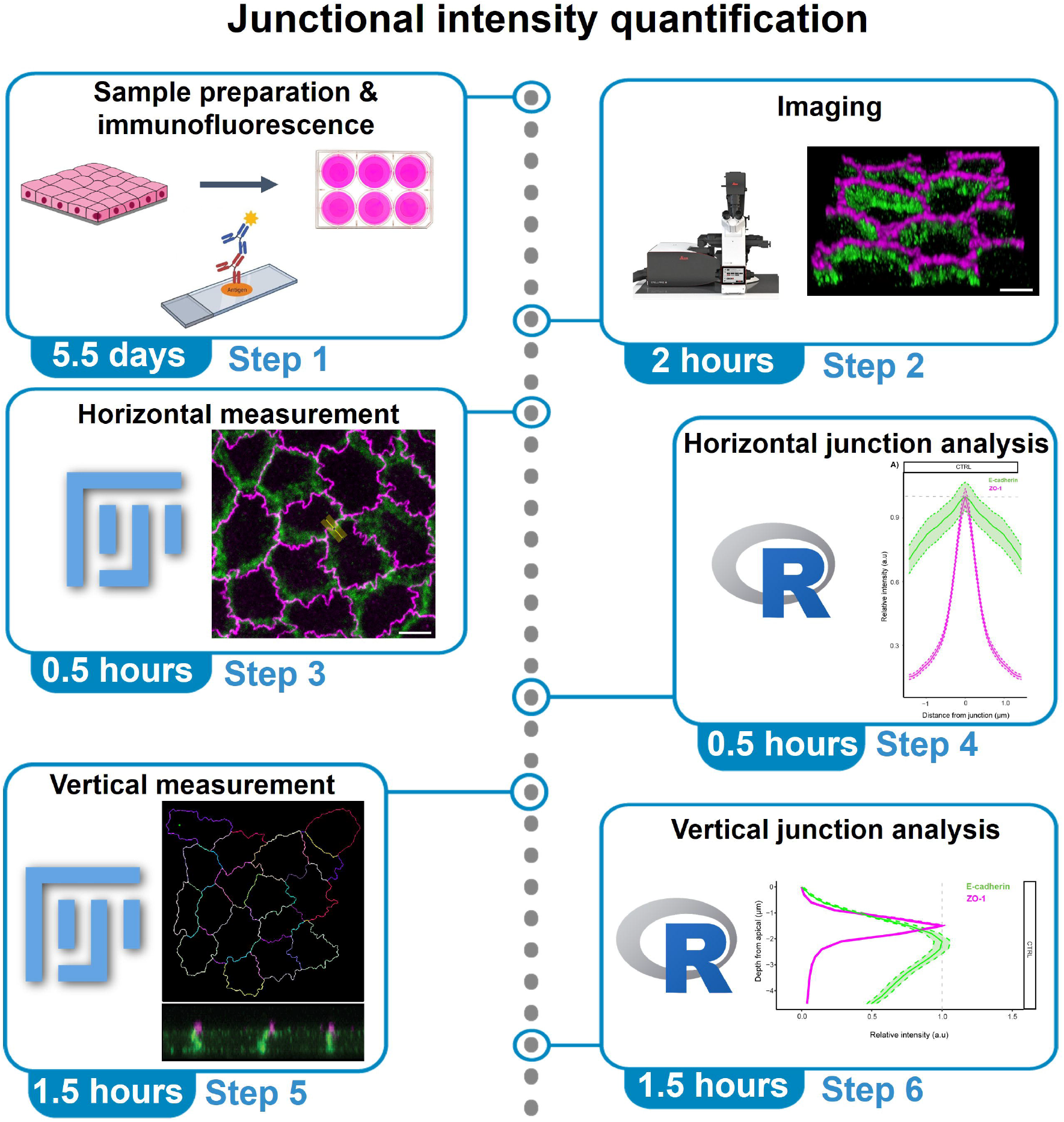

## Before you begin

In before you begin steps 1-8, as well as step-by-step method details 1-12 we outline the protocol for measuring the distribution of AJ (E-cadherin) and TJ (ZO-1) proteins across mature cell-cell junctions in MDCK cells. These fluorescence intensity measurements are obtained using Fiji/ImageJ by defining multiple regions of interest (ROIs) as lines perpendicular to the junctions within an acquired field of view. Intensity profiles are then measured across the entire length of each ROI, allowing for efficient quantification of a large number of junctions within a monolayer. Instructions for compiling and graphing this data using RStudio are provided in step-by-step method details 13-25.

In before you begin steps 1-8, as well as step-by-step method details 26-52, we describe how to measure the distribution of AJ and TJ proteins along the Z-axis of the junctions. Individual cell junctions are defined through segmentation in Tissue Analyzer^2^, with the resulting segmentation exported as regions of interest (ROIs) to ImageJ. When paired with the RStudio instructions for data compilation and visualization found in step-by-step method details 53-65, this method is an efficient approach for characterizing single junctions on a large scale.

Here we use E-cadherin and ZO-1 as examples of AJ and TJ proteins respectively, but these techniques can be adapted to other junctional protein markers of interest. Because these protocols rely on quantitative microscopy image analysis, maintaining similar acquisition parameters across different samples and conditions is essential for reproducibility.

### Sample preparation

#### Timing: 5 days

1. Culture MDCK cells in DMEM media supplemented with 10 % fetal bovine serum (FBS) and 1x Penicillin-Streptomycin.
2. Split 600,000 MDCK cells into a 6 well plate (35 mm) containing 12 mm glass coverslips (#1.5).
3. Incubate the cells for four days to allow the formation of a polarized monolayer with mature cell-cell junctions. Replace the media every other day, or more frequently as needed to maintain cell health.
4. Follow the immunofluorescence protocol described in Awadia et al. (2019)^3^, staining for the desired junctional proteins. **Critical:** Include a at least one marker that is known to be strongly and specifically localized to cell-cell junctions, such as ZO-1, for ease of automated segmentation in later steps.

### Image acquisition

#### Timing: 1-3 h

5. Use a high-numerical-aperture 63x oil immersion objective.
6. Apply a small drop of immersion oil to the surface of each coverslip on the slide. Invert the slide so that the immersion oil on the selected coverslip contacts the objective. **Critical:** Use a low-autofluorescence immersion oil specifically designed for the objective being used.
7. Visualize the stained junctional proteins using fluorescence microscopy and acquire at least five random fields of view for each condition. Make sure to acquire enough z-planes to encompass the entire apicobasal distribution of the proteins in the Z axis of the monolayer, as well as a few additional slices above and below the monolayer. **Critical**: It is essential to maintain identical acquisition settings across and within conditions to ensure pixel intensity values are comparable.
8. Save the raw microscopy images in a directory that can be accessed during quantification steps.

## Key resources table

| REAGENT or RESOURCE | SOURCE | IDENTIFIER |
| --- | --- | --- |
| **Antibodies** |  |  |
| Anti-E-cadherin (Rabbit) | Cell Signaling Technology | Cat#24E10 |
| Anti-ZO-1 (Mouse) | Thermo Fisher Scientific | Cat#339100 |
| Secondary Alexa-594 (Anti-Mouse) | Thermo Fisher Scientific | Cat#A11005 |
| Secondary Alexa-488 (Anti-Rabbit) | Thermo Fisher Scientific | Cat#A11008 |
| **Chemicals, peptides, and recombinant proteins** |  |  |
| Formaldehyde solution 4%, buffered, pH 6.9 | Sigma-Aldrich | Cat#1004960700 |
| ProLong™ Diamond Antifade Mountant | Thermo Fisher Scientific | Cat#P36961 |
| Hoechst 34580 | Sigma-Aldrich | Cat#63493 |
| **Experimental models: Cell lines** |  |  |
| MDCKII T23 | Macara Lab |  |
| **Software and algorithms** |  |  |
| Fiji/ImageJ | Available from NIH | https://imagej.net/software/fiji/ |
| RStudio version 1.4.1106 | RStudio | https://www.rstudio.com |
| LAS X | Leica | https://www.leica-microsystems.com/products/microscope-software/p/leica-las-x-ls/downloads/ |
| **Other** |  |  |
| Coverslips, 12 mm, #1.5 | Fisher Scientific | Cat#48382-063 |
| Glass slides | Fisher Scientific | Cat#10118-596 |
| Leica laser scanning confocal system | Leica | Stellaris 5 |

## Materials and equipment

This protocol was implemented using a Leica Stellaris 5 laser scanning confocal microscope, equipped with an HC PL APO 63X/1.40 OIL CS2 objective, hybrid (HyD) detectors, and LAS X software. The system requirements for this protocol are highly flexible. Any confocal fluorescence microscope capable of acquiring complete three-dimensional images of the cell-cell junctions is appropriate for this pipeline.

### Step-by-step method details

For horizontal profiling of cell-cell junctions, follow steps 1-25. For vertical profiling of the junctions, follow steps 26-65.

### Horizontal profiling: Junction quantification with Fiji/ImageJ

#### Timing: 30 minutes

The goal of this analysis is to understand the distribution of different proteins across an epithelial junction or changes in the distribution of the same protein in different conditions. Usually, a marker that defines junctions is used (ZO-1) and the distribution of a second protein is analyzed (E-cadherin). When two different experimental conditions are compared, like CTRL and KD, it is essential to follow a consistent naming convention for image files. The image from the control condition should be named with the prefix “CTRL_” (including the underscore), while the naming format for the other condition is flexible. If you are analyzing multiple fields of view (FOV) per condition it is important to indicate this in the name of the image as follow: “CTRL_FOV1_” or “KD_FOV1_”. This naming system ensures proper analysis of the data in R Studio. If you are not comparing two conditions or different FOV, it is not necessary to follow this naming format.

1. Open the image to be quantified by dragging it into FIJI/ImageJ or by using File>Open (Figure 1A).
2. Create a new macro in Fiji/ImageJ by going to File > New > Script. **Note:** If the macro is already saved, open it by dragging the macro file into the taskbar in Fiji/ImageJ, and skip to step 6.
3. Define the coding language in the macro window by navigating to Language > ImageJ Macro.
4. Copy the following code into the script editor (Figure 1B):

**Figure 1:**
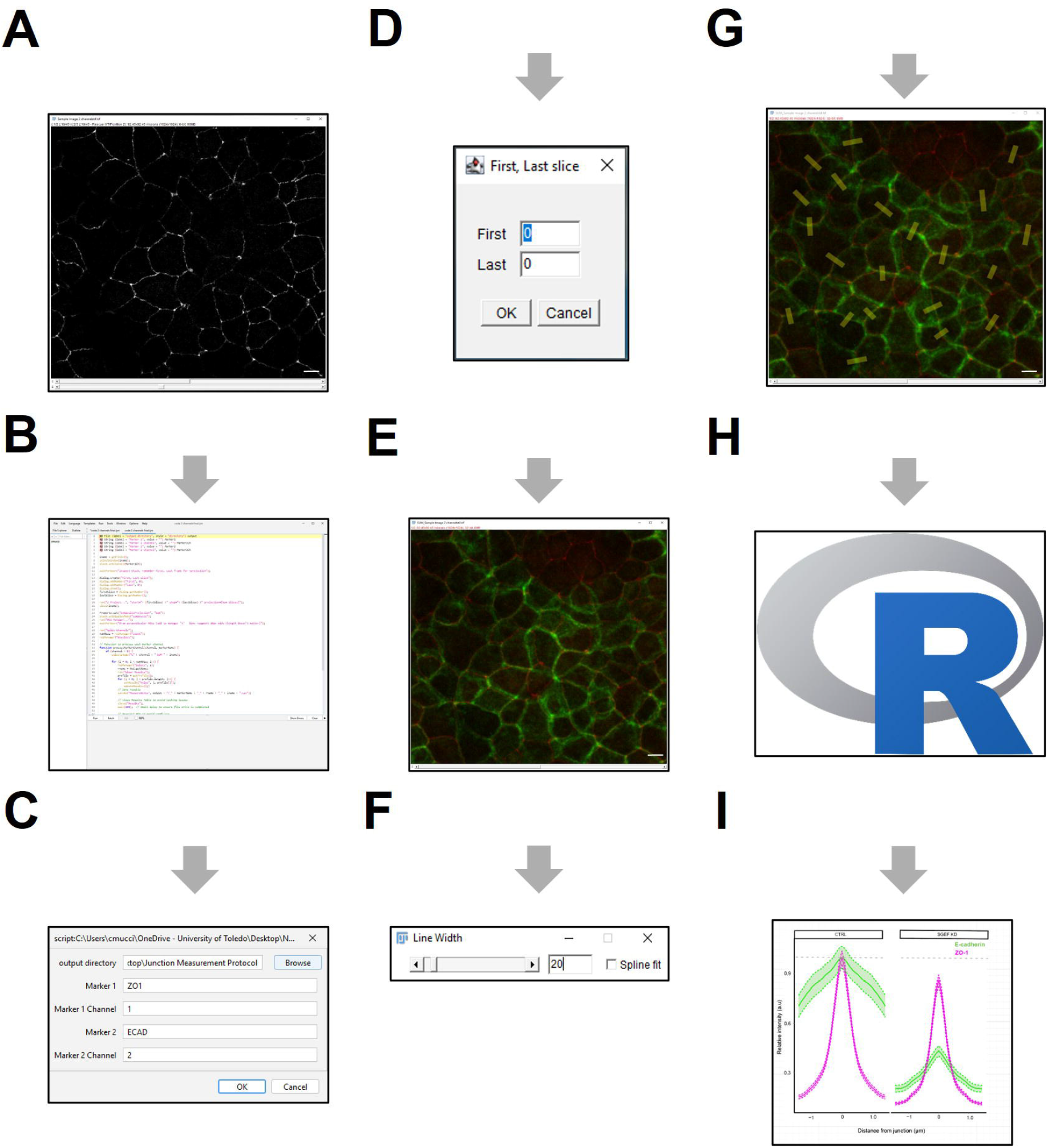
Horizontal profiling quantification pipeline (A-F) Workflow of image analysis used to quantify junctions horizontally.

~~~
>#@ File (label = “output directory”, style = “directory”) output
>#@ String (label = “Marker 1”, value = ““) Marker1
>#@ String (label = “Marker 1 Channel”, value = ““) Marker1Ch
>#@ String (label = “Marker 2”, value = ““) Marker2
>#@ String (label = “Marker 2 Channel”, value = ““) Marker2Ch
>iname = getTitle();
>selectWindow(iname);
>Stack.setChannel(Marker1Ch);
>waitForUser(“Inspect Stack, remember First, Last frame for >projection”);
>Dialog.create(“First, Last slice”);
>Dialog.addNumber(“First”, 0);
>Dialog.addNumber(“Last”, 0);
>Dialog.show();
>firstSlice = Dialog.getNumber();
>lastSlice = Dialog.getNumber();
>run(“Z Project…”, “start=“+ (firstSlice) +” stop=“+ (lastSlice) +” projection=[Sum Slices]”);
>close(iname);
>Property.set(“CompositeProjection”, “Sum”);
>Stack.setDisplayMode(“composite”);
>run(“ROI Manager…”);
>waitForUser(“Draw perpendicular ROIs (add to manager ‘t’ - line >segment 20px wide (length doesn’t matter)”);
>run(“Split Channels”);
>numROIs = roiManager(“count”);
>roiManager(“Deselect”);
>// Function to process each marker channel
>function processMarkerChannel(channel, markerName) {
>  if (channel > 0) {
>    selectImage(“C” + channel + “-SUM_” + iname);
>    for (i = 0; i < numROIs; i++) {
>      roiManager(“Select”, i);
>      rname = Roi.getName;
>      run(“Clear Results”);
>      profile = getProfile();
>      for (j = 0; j < profile.length; j++) {
>         setResult(“Value”, j, profile[j]);
>         updateResults();}
>      // Save results
> saveAs(“Measurements”, output + “/_” + markerName + “_” +
rname + “_” + iname + “.csv”);
>      // Close Results Table to avoid locking issues
>      close(“Results”);
>      wait(100); // Small delay to ensure file write is completed
>      // Deselect ROI to avoid conflicts
>      roiManager(“Deselect”);
>    run(“Collect Garbage”);}}}
>// Process Marker 1
>processMarkerChannel(Marker1Ch, Marker1);
>// Process Marker 2
>processMarkerChannel(Marker2Ch, Marker2);
>// Close all images
>run(“Close All”);
>// Close ROI Manager
>if (isOpen(“ROI Manager”)) {
>  roiManager(“reset”);
>  run(“ROI Manager…”);
>  close(“ROI Manager”);}
>// Final garbage collection
>run(“Collect Garbage”);
>print(“⍰ Processing complete. All files saved and windows closed.”);
~~~

5. Run the script by clicking the “Run” button at the bottom of the macro screen **Note:** User comments are indicated by “//” in the code and will not interfere with the execution.
6. A prompt window will appear in Fiji/ImageJ, allowing the user to specify destination folder for the quantification files. Additionally, the user will designate the specific channels in the original image to be quantified (Figure 1C). The order of the channels can be found with the channel (C) slider in the original image. **Note:** The name assigned by the user to each channel will be used by ImageJ to save the corresponding files.
7. Next, a pop-up window asks the user to inspect the stack and identify the first and last slice with signal. To do that, select the channel with a cell marker like ZO-1 to identify the first and last slice to be used for quantification.
8. Add the numbers for the first and last slice in the following pop-up window (Figure 1D).
9. Next, the code will integrate all Z slices into a single plane using the sum slices option and generate a composite image (Figure 1E).
10. A new pop-up window is going to show up asking the user to draw ROIs perpendicular to the junctions to be measured. There are no limitations on the numbers of ROIs that can be drawn. The number will be determined by the user.
11. Double click in the line segment tool and change the line segment to 20 pixels (Figure 1F), then draw perpendicular ROIs on the junctions to be measured and add them to the ROI manager with shortcut “t”. The length of the ROIs should be at least 32 pixels long, one ROI per junction. **Note:** Depending on where the ROI is located within the junction, the measurements can be different. For example, one ROI placed at the center of the junction vs. another ROI placed near the tricellular junction. The placement of the ROI should be determined based on your specific biological question.
12. Once all the ROIs have been drawn (Figure 1G), click “OK”. ImageJ will then measure the fluorescent intensity of each ROI in both channels. When finished, the results will be saved as a .CSV file in the previously selected folder (Step 6), and all open windows will be closed in ImageJ. Each ROI for each channel is saved as an individual file. This is important for the subsequent analysis in R Studio (Steps 13-25). **Note:** Sample datasets generated from the Fiji/ImageJ horizontal junction quantifications are provided in Table 1 and 2. As mentioned above, the code saves each ROI for each channel as individual files (Steps 1-12), but we compiled multiple ROIs in the table as an example.

**Table 1:** Example of raw ECAD intensity data from horizontal junction quantification in Fiji/ImageJ. Only for this example table, all the ROI are in the same table. After Fiji/ImageJ quantification you will have one CSV file per ROI.

| ROI 1 |  | ROI 2 |  | ROI 3 |  |
| --- | --- | --- | --- | --- | --- |
| Slice | Value | Slice | Value | Slice | Value |
| 1 | 32.623 | 1 | 20.318 | 1 | 21.487 |
| 2 | 33.311 | 2 | 21.628 | 2 | 24.338 |
| 3 | 36.168 | 3 | 21.015 | 3 | 24.199 |
| 4 | 33.172 | 4 | 18.431 | 4 | 25.352 |
| 5 | 30.801 | 5 | 18.598 | 5 | 25.475 |
| 6 | 33.321 | 6 | 17.956 | 6 | 25.446 |
| 7 | 30.5 | 7 | 17.077 | 7 | 25.715 |
| 8 | 28.745 | 8 | 18.775 | 8 | 27.061 |
| 9 | 29.673 | 9 | 18.341 | 9 | 27.766 |
| 10 | 30.217 | 10 | 19.407 | 10 | 29.185 |
| 11 | 34.452 | 11 | 19.791 | 11 | 32.252 |
| 12 | 39.72 | 12 | 20.019 | 12 | 32.842 |
| 13 | 45.542 | 13 | 21.14 | 13 | 33.343 |
| 14 | 51.217 | 14 | 22.393 | 14 | 33.793 |
| 15 | 56.642 | 15 | 26.198 | 15 | 34.572 |
| 16 | 62.775 | 16 | 30.669 | 16 | 37.499 |
| 17 | 73.289 | 17 | 35.56 | 17 | 39.917 |
| 18 | 86.067 | 18 | 38.908 | 18 | 42.093 |
| 19 | 92.809 | 19 | 38.073 | 19 | 48.098 |
| 20 | 96.45 | 20 | 42.8 | 20 | 50.279 |
| 21 | 97.663 | 21 | 43.84 | 21 | 50.647 |
| 22 | 91.34 | 22 | 41.441 | 22 | 48.928 |
| 23 | 86.281 | 23 | 36.219 | 23 | 49.355 |
| 24 | 81.428 | 24 | 31.429 | 24 | 43.977 |
| 25 | 78.049 | 25 | 27.372 | 25 | 35.932 |
| 26 | 73.562 | 26 | 24.75 | 26 | 30.208 |
| 27 | 73.746 | 27 | 24.598 | 27 | 25.166 |
| 28 | 73.266 | 28 | 27.942 | 28 | 23.735 |
| 29 | 70.63 | 29 | 26.203 | 29 | 21.504 |
| 30 | 63.61 | 30 | 26.154 | 30 | 20.547 |
| 31 | 56.385 | 31 | 26.229 | 31 | 22.377 |
| 32 | 54.306 | 32 | 26.014 | 32 | 20.959 |
| 33 | 47.864 | 33 | 27.041 | 33 | 19.663 |
| 34 | 43.482 | 34 | 28.743 | 34 | 18.838 |
| 35 | 43.865 | 35 | 28.272 | 35 | 19.619 |
| 36 | 40.52 | 36 | 28.498 | 36 | 18.626 |
| 37 | 35.787 | 37 | 29.08 | 37 | 19.11 |
| 38 | 32.693 | 38 | 27.826 | 38 | 20.197 |
| 39 | 34.988 | 39 | 26.86 | 39 | 19.945 |
| 40 | 37.801 | 40 | 26.695 | 40 | 21.393 |
| 41 | 37.759 | 41 | 25.436 | 41 | 20.006 |

**Table 2:** Example of raw ZO1 intensity data from horizontal junction quantification in Fiji/ImageJ. Only for this example table, all the ROI are in the same table. After Fiji/ImageJ quantification you will have one CSV file per ROI.

| ROI 1 |  | ROI 2 |  | ROI 3 |  |
| --- | --- | --- | --- | --- | --- |
| Slice | Value | Slice | Value | Slice | Value |
| 1 | 13.507 | 1 | 14.931 | 1 | 11.816 |
| 2 | 13.471 | 2 | 14.248 | 2 | 12.999 |
| 3 | 15.113 | 3 | 14.143 | 3 | 12.493 |
| 4 | 13.109 | 4 | 13.715 | 4 | 14.754 |
| 5 | 13.609 | 5 | 13.863 | 5 | 15.705 |
| 6 | 14.167 | 6 | 13.888 | 6 | 15.984 |
| 7 | 14.408 | 7 | 11.779 | 7 | 16.833 |
| 8 | 12.589 | 8 | 13.855 | 8 | 18.042 |
| 9 | 12.543 | 9 | 13.696 | 9 | 16.412 |
| 10 | 14.615 | 10 | 12.469 | 10 | 16.423 |
| 11 | 15.668 | 11 | 12.408 | 11 | 16.735 |
| 12 | 17.543 | 12 | 12.81 | 12 | 18.935 |
| 13 | 18.822 | 13 | 13.885 | 13 | 16.868 |
| 14 | 18.498 | 14 | 13.25 | 14 | 17.773 |
| 15 | 20.704 | 15 | 14.462 | 15 | 19.883 |
| 16 | 20.647 | 16 | 13.805 | 16 | 21.128 |
| 17 | 19.879 | 17 | 15.09 | 17 | 21.254 |
| 18 | 19.23 | 18 | 16.574 | 18 | 24.729 |
| 19 | 22.206 | 19 | 19.613 | 19 | 29.517 |
| 20 | 22.109 | 20 | 20.355 | 20 | 36.23 |
| 21 | 23.348 | 21 | 21.777 | 21 | 42.201 |
| 22 | 21.458 | 22 | 21.731 | 22 | 41.959 |
| 23 | 19.125 | 23 | 20.784 | 23 | 38.488 |
| 24 | 16.446 | 24 | 19.439 | 24 | 32.586 |
| 25 | 15.944 | 25 | 18.033 | 25 | 28.719 |
| 26 | 16.214 | 26 | 17.783 | 26 | 24.512 |
| 27 | 15.541 | 27 | 15.888 | 27 | 19.472 |
| 28 | 15.882 | 28 | 17.871 | 28 | 17.876 |
| 29 | 14.908 | 29 | 19.061 | 29 | 15.786 |
| 30 | 16.56 | 30 | 18.917 | 30 | 14.643 |
| 31 | 15.492 | 31 | 18.589 | 31 | 13.393 |
| 32 | 14.724 | 32 | 18.468 | 32 | 12.766 |
| 33 | 14.042 | 33 | 16.821 | 33 | 12.26 |
| 34 | 13.421 | 34 | 15.084 | 34 | 12.239 |
| 35 | 13.483 | 35 | 14.181 | 35 | 11.993 |
| 36 | 12.201 | 36 | 13.456 | 36 | 13.778 |
| 37 | 13.081 | 37 | 14.986 | 37 | 13.285 |
| 38 | 13.54 | 38 | 16.608 | 38 | 12.935 |
| 39 | 14.424 | 39 | 17.053 | 39 | 13.668 |
| 40 | 15.884 | 40 | 15.72 | 40 | 13.474 |
| 41 | 15.062 | 41 | 15.648 | 41 | 12.388 |

### Horizontal profiling: Data Analysis with RStudio

#### Timing: 10 minutes

The layout of RStudio differs from that of the Fiji/ImageJ macro window. In RStudio, the Source pane (typically in the upper left) is where code is written, edited, and executed. The Console, located in the bottom left, displays real-time execution of commands and maintains a history of previously run code within the session. Any errors that occur will be shown in this window. A “>” prompt in the Console indicates that the code has executed successfully, and that R is ready for the next command. The Environment pane, found in the upper right, contains active dataframes created during the session, which can be viewed at any time by clicking on them. Lastly, the Plots pane in the bottom right displays graphs generated by the code.

13. Download and install the R programming language along with the RStudio software interface.
14. Open RStudio and create a new script by navigating to File > New File > R Script. Save the script in the same directory as the ZO-1 and E-cadherin CSV files. **Note:** To run the RStudio code line by line, use the “Ctrl + Enter” keys. User comments are indicated by “#” in the code and will not interfere with execution.
15. Use the following code to load the necessary R packages:

~~~
pacman::p_load(stringr,rstudioapi,tidyverse, wesanderson, Rmisc, ggplot2, plotly, carData, car, inferr, agricolae, multcompView, dplyr, plyr, multcomp, corrplot, gridExtra, grid, ggpubr, ClusterRankTest, readr,viridis, ggforce, purrr)
~~~

**Note:** The packages need to be loaded only once per RStudio session. Each package has specific functions and syntax for data analysis.

16. Set the working directory to the location of the script using the following commands:

~~~
setwd(dirname(rstudioapi::getActiveDocumentContext()$path))
dir<-getwd()
~~~

17. Identify files by name structure with the following code: **Note:** in the (“ZO1*.csv”) and (“ECAD*.csv”) part of the code, the name MUST match the name in the .CSV file (defined in step 6), including capital letters, if not, the code will not work.

~~~
>filenameZO1 <-list.files(path = dir, pattern = glob2rx(“_ZO1_*.csv”), full.names = FALSE, recursive = TRUE, include.dirs = FALSE) #full.names FALSE, although counter intuitive, it means you save the name and not the path
>my_dataZO1 <-list.files(path = dir, pattern = glob2rx(“_ZO1_*.csv”), full.names = TRUE, recursive = TRUE) %>% #reading the path to your CSV files
>map_dfr(read_csv, .id = “path”) %>% #mapping the “path” of each file to the individual rows
>group_by(path) %>% #grouping by path (or file)
>nest() %>% #nesting into a new matrix that will have the same dimensions as your filenames list
>cbind(filenameZO1)%>% #binding the list names as a column to the nested file
>unnest(c(data))%>% #unnesting (each datapoint in the nested row will copyhave the file name)
>dplyr::rename(ZO1 = Value)
>filenameEcad <-list.files(path = dir, pattern = glob2rx(“_ECAD_*.csv”), full.names = FALSE, recursive = TRUE, include.dirs = FALSE) #full.names FALSE, although counter intuitive, it means you save the name and not the path
>my_dataEcad <-list.files(path = dir, pattern = glob2rx(“_ECAD_*.csv”), full.names = TRUE, recursive = TRUE) %>% #reading the path to your CSV files
>map_dfr(read_csv, .id = “path”) %>% #mapping the “path” of each file to the individual rows
>group_by(path) %>% #grouping by path (or file)
>nest() %>% #nesting into a new matrix that will have the same dimensions as your filenames list
>cbind(filenameEcad)%>% #binding the list names as a column to the nested file
>unnest(c(data))%>% #unnesting (each datapoint in the nested row will copyhave the file name)
>dplyr::rename(Ecad = Value)
~~~

18. Create a unique dataframe called “my_data” binding all columns with the following code:

~~~
>my_data <-my_dataZO1 %>% cbind(my_dataEcad) %>%
>dplyr::select(…4, ZO1, Ecad)%>%
>separate(…4, c(NA, NA, “ROI”, “MUT”, “FOV”), sep=“_”) #Extracts
meta data from File name, important to structure files always the same
~~~

19. Run the following code to filter the data and keep the ROIs that are at least 3 µm long. This code is going to pick the brightest point of ZO-1 in each ROI and 1.442768 µm to each side of the brightest point. The number 16 corresponds to the number of pixels that are in 1.442768 µm. This number will vary depending on your microscopy system and with your acquisition parameters and may need to be changed accordingly. **Note:** If you want less resolution, you can decrease the number of pixels. To do that, you will need to replace the number 16 in the lines of codes below and you will also need to replace the numbers in the lines of code in step 20. If your pixel resolution is 0.090173 µm you will only need to modify the number “1.442768” that is the result of multiplying the number of pixels by the resolution.

~~~
> filtered <-my_data %>% group_by(ROI, FOV, MUT) %>%
> mutate(maxZO1 = which.max(ZO1), ##Added this lines of code to discard any ROI that doesn’t fullfill the Slice. This means the user drew a short ROI, and is not at least 3um long.
>lastZO1 = n()-maxZO1)%>%
>filter(maxZO1 >= 16) %>%
>filter(lastZO1 >= 16)%>%
>slice((which.max(ZO1)-16) : (which.max(ZO1) +16))%>% #16pixels because its roughly 1.5um to each side of the brightest point, If used doesnt want this much resolution they can decrease the number >mutate(id = as.numeric(row_number()-1))
~~~

20. Create a positive and negative dataframe to center the data around the brightest pixel (labeled as 0). Positive dataframe will go from 0 to 1.44 µm and negative dataframe will go from –1.44 to 0 µm. **Note:** The ID is converted to microns using the pixel depth of the image, which may vary depending on acquisition parameters.

~~~
>positivedataframe <-filtered %>% mutate(centered = (id * 0.090173)-
1.442768)%>% ## 0.09 is µm resolution, 1.44 is just 16 pixels times
0.09, this way the brightest pixel will be labeled as 0
 mutate(centered = round(centered, 2))
>inverteddataframe <-filtered %>% mutate(centered = (id * −0.090173
)+1.442768) %>%
mutate(centered = round(centered, 2))
~~~

21. Create a dataframe using the following code:

~~~
> dataframe<-positivedataframe %>% bind_rows(inverteddataframe)%>%
> pivot_longer(“ZO1”:”Ecad”, names_to = “Signal”, values_to = “Intensity”)%>%
> ungroup()
> summary <-dataframe %>% group_by(MUT, Signal, centered)%>%
> summarise(Mean=mean(Intensity), Min=min(Intensity))
>             #gets maximum and minimum values from previously defined Signals,
if you have more stainings/FP add by copy and pasting accordingly
> MaxEcad <-summary %>% group_by(Signal, MUT)%>%
> summarise(Max=max(Mean), Min=min(Min))%>%
> filter (MUT==“CTRL” & Signal==“Ecad”)
>     MaxZO1 <-summary %>% group_by(Signal, MUT)%>%
>      summarise(Max=max(Mean),Min=min(Min)) %>%
>     filter(MUT==“CTRL” & Signal==“ZO1”)
~~~

22. Normalization of the data to make both experimental conditions comparable.

~~~
> dataframe <-dataframe %>% mutate(Intensity = case_when(
           Signal==“ZO1” ~ (Intensity - MaxZO1$Min)/( MaxZO1$Max -
MaxZO1$Min),
          Signal==“Ecad” ~ (Intensity - MaxEcad$Min)/( MaxEcad$Max
- MaxEcad$Min)))
~~~

23. Create a dataframe mean.

~~~
> dataframeMean <- dataframe %>% group_by(MUT, Signal, centered) %>%
> summarise(sd = sd(Intensity),
                 n = n(), # Calculate sample size
                 SE = sd / sqrt(n), # Compute Standard Error
                 Mean = mean(Intensity))
~~~

24. Create the graph (Figure 1I). This code is going to plot all the measuraments from all the channels in the same graph. If you want a separate graph for each channel add this piece of code: rows=vars (Signal) to the line “facet_grid”. The full code will be facet_grid(cols=vars (MUT), rows=vars (Signal)).

~~~
> plot <-ggplot(dataframeMean, aes(x=centered, y=Mean)) + ##general
data aes-what it would be graphed
> facet_grid(cols=vars(MUT))+ geom_path(aes(color=Signal), size=1)+
geom_ribbon(aes(fill=Signal, ymax = Mean+SE, ymin = Mean-SE),alpha=0)+
geom_line(aes(y = Mean + SE, color=Signal), linetype=“dashed”, size=0.5)
+ # Upper SE line geom_line(aes(y = Mean - SE, color=Signal),
linetype=“dashed”, size=0.5) + # Lower SE line theme_classic()+
 scale_shape_manual(values=c(3,4,8,15,16,17,18,19))+
 scale_color_manual(values=c(“green”,”magenta”))+ #scale_fill_manual(values=c(“green”, “magenta”)) +
 xlab(“Distance from junction (µm)”)+
 ylab(“Protein Intensity (a.u)”)+ guides(shape=FALSE)+
guides(color=FALSE)+
 guides(fill=FALSE)
>plot
~~~

25. In the Plots panel, navigate to Export > Save as Image, and save the graph to the desired location.

### Vertical profiling: Image preparation for segmentation

#### Timing: 5 minutes

Here, we outline the steps for preparing raw image files of any format for segmentation of individual junctions using Fiji/ImageJ.

26. Open the raw image to be quantified in Fiji/ImageJ and save a copy in the TIFF file format to preserve the quality of the original data: File > Save as > Tiff, including all Z slices and channels. Save the TIFF file in a folder that is dedicated only to this quantification. **Critical:** In the RStudio portion of this protocol, quantified values are categorized based on experimental variables (e.g., “Experiment” or “Condition”) so that the code may be adapted for large-scale data compilation. To allow automated categorization, file names must be structured with underscores separating these key identifiers (e.g., Exp1_CTRL_.tif).
27. Isolation of the segmentation channel: To accurately segment cells within a monolayer, the channel most strongly localized to the cell-cell junctions should be selected.
  a. Duplicate the image (Image>Duplicate), keeping all Z slices, and choose only the desired segmentation channel in the prompt window (Figure 2A). **Note:** In this protocol, we use the ZO-1 channel to define the tight junctions that connect the cells. However, any marker that strongly localizes to the cell boundaries can be used.
28. Max projection of Z slices: Because the junctional protein of choice may localize in slightly different planes across the monolayer, all Z slices should be compressed into a single plane using a max intensity projection.
  a. To create a max projection, run the following Fiji/ImageJ function: Image > Stacks > Z project, and select “Max Intensity” from the Projection Type drop-down menu (Figure 2B). **Note:** If the ZO-1 signal is dim in the KD or other conditions, the brightness can be adjusted in Fiji/ImageJ: Image > Adjust > Brightness/Contrast. However, the brightness should not be altered in the image saved in step-by-step methods details step 26.
29. Save the new max projection image in the TIFF file format. While the file name can be arbitrary, using a concise name is recommended for easy reference in following steps.

**Figure 2:**
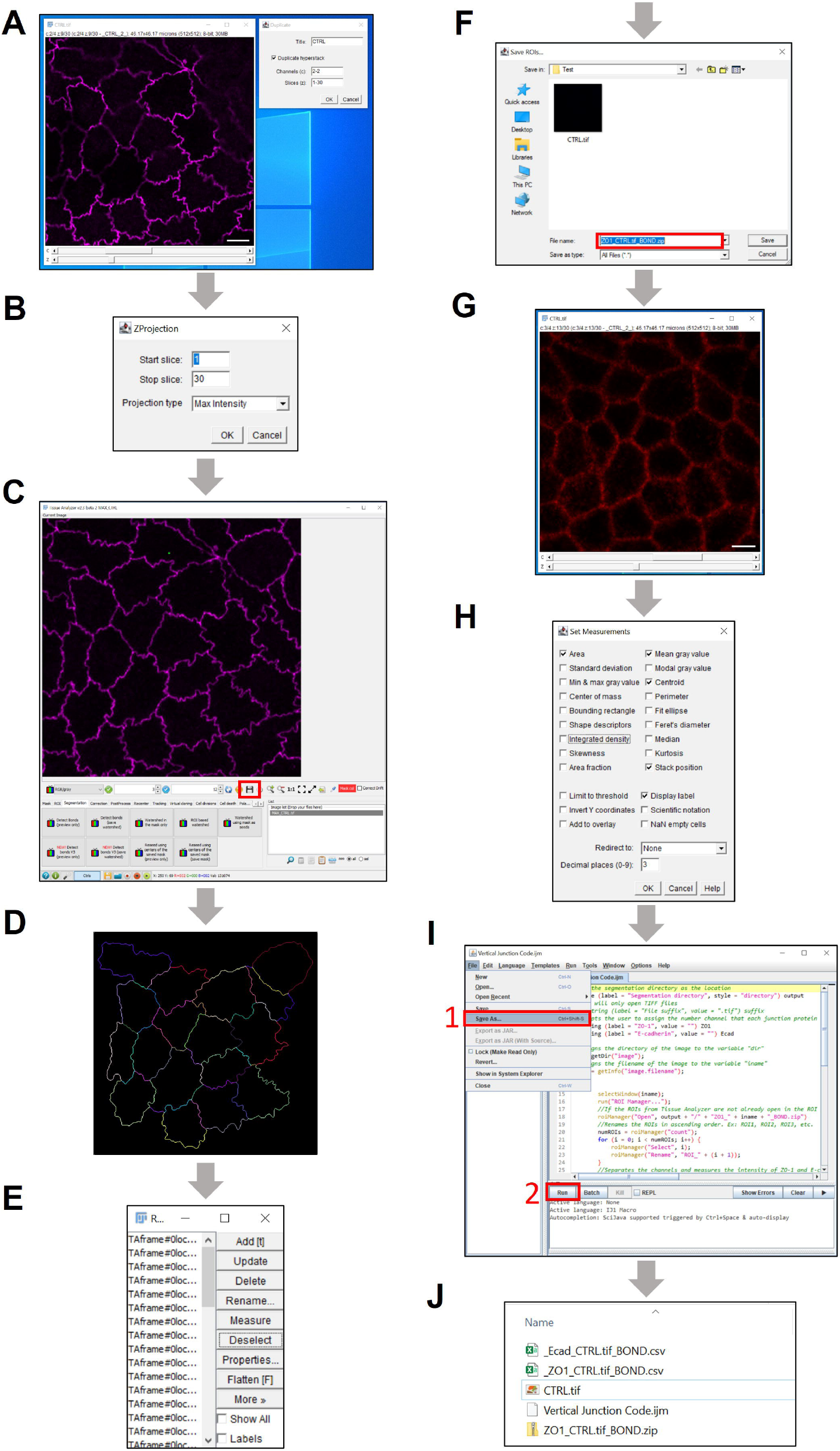
Vertical profiling quantification pipeline. (A)Segmentation channel duplication in Fiji/ImageJ. Scale bar 5 µm. (B)Z-stack maximum intensity projection in Fiji/ImageJ. (C)Tissue Analyzer layout and selection of max projection in the software. “Save” icon is indicated by a red box. (D)Finalized junction segmentation in Tissue Analyzer. (E)Exported ROI list in Fiji/ImageJ ROI Manager. (F)ROI zip file save location. (G)TIFF image with all channels and z slices. Scale bar 5 µm. (H)Fiji/ImageJ measurement parameters pop-up window. (I)Saving and running the Fiji/ImageJ macro. (J)Final output in the quantification folder from execution of the Vertical Junction Code Fiji/ImageJ macro.

### Vertical profiling: Segmentation with Tissue Analyzer

#### Timing: 20 minutes

Here we describe an automated method for segmenting the individual cells within the max-projection image obtained in the previous step. Although manual segmentation using the drawing tool in Fiji/ImageJ is possible, it is time-consuming and may yield results that are more subjective. The following workflow outlines the steps for cell segmentation using the Tissue Analyzer plugin in Fiji/ImageJ.

**Note:** Any previously established segmentation method may be used if it allows the individual junctions to be exported to Fiji/ImageJ as regions of interest (ROIs).

30. Download the Tissue Analyzer plugin: Navigate to Help > Update > Manage Update Sites > select Tissue Analyzer > Apply Changes. **Note:** If the Tissue Analyzer plugin is already installed, skip to Step 32.
31. Close and restart Fiji/ImageJ.
32. Open Tissue Analyzer: Go to Plugins > Tissue Analyzer.
33. Drag the max projection TIFF file from step 29 into the image list in Tissue Analyzer and select it (Figure 2C).
34. Under the Segmentation tab click “Detect bonds V3 (preview only)” to configure the segmentation parameters.
  a. Uncheck the Optional PreProcess box
  b. For segmentation using ZO-1 or a similar stain, set a Strong blur of 4.5 and a Weak blur of 1. c.
  c. Keep all other settings as default: “compare to kuwahara pass,” “compare to max pass,” and “finalize mask” should be selected.
  d. Click OK to apply the parameters and view the preview segmentation.
  e. Adjust the Strong blur setting as needed for optimal results. **Note:** The automated segmentation may not be fully accurate, so some manual corrections should be anticipated in subsequent steps.
35. Once the desired parameters have been set, select “Detect bonds V3 (save watershed)”, using the same settings that were optimized in the previous step.
36. Manual correction: To refine segmentation, create and delete bonds as follows:
  a. Add bonds: Hold down the left mouse button, trace on top of the junction, and press enter.
  b. Delete bonds: Hold down the right mouse button, drag over unwanted bonds, and press enter. A short line across the bond is sufficient for deletion, it does not need to fully cover the junction.
  c. Save the corrections using the icon in the lower right corner before proceeding (Figure 2C, red box).
  **Critical:** Delete all junctions that contact the image border at this stage, because attempting this later will be considerably more tedious.
37. Navigate to the PostProcess tab and select “Finish all.” Begin with the default settings and adjust as needed. “Fix mask outlier pixels (Recommended)” should be selected. **Note:** A lower “4-way vertex vs bond cut off” value may better preserve junction curvature.
38. Select “Check finish all.”
39. Go to the Tracking tab and select “Track cells (static tissue)” with the default settings:
  a. Max Displacement in the Displacement exclusion criteria should be 25.
  b. “No” should be selected for “Exclude biggest cell”.
40. Click “Check track”.
41. Under the Tracked bonds tab, select “Track bonds”.
42. Click “Check track bonds” to finalize individual junction segmentation (Figure 2D). **Note:** If the initial segmentation is modified, update the Tracking and Tracked bonds masks using the “Update track mask” and “Update track bonds” options respectively.
43. In the Fiji/ImageJ taskbar, double-click the Line Segment Tool and confirm that the line width is set to 1 pixel before proceeding.
44. Select “Convert TA bonds to IJ ROIs” and click OK in the pop-up window. This will export each segmented junction to the Fiji/ImageJ ROI Manager as a unique ROI (Figure 2E).

### Vertical profiling: Junction quantification with Fiji/ImageJ

#### Timing: 20 minutes

Here, we describe a method using the Fiji/ImageJ macro function to apply the ROIs that were previously generated in Tissue Analyzer for profiling signal intensity at each individual junction across the Z-stack using the original image. The output of this procedure is a .CSV file for each junction marker that contains the acquired measurements.

45. Save all ROIs: In the ROI manager in Fiji/ImageJ, ensure all ROIs are deselected by clicking “Deselect”. Then, navigate to More > Save.
  a. Name the **ROI zip file** in the format “ZO1_imagefilename.tif_BOND.zip”, where imagefilename corresponds to **the name** of the entire TIFF image from step-by-step method details, step 26 (e.g., ZO1_CTRL.tif_BOND.zip).
  b. Save the ROI zip file in the quantification folder (Figure 2F).
46. Open the TIFF image saved in step-by-step method details, step 26 by dragging it to the Fiji/ImageJ taskbar (Figure 2G). This TIFF file should include all channels and Z slices acquired from the original image.
47. Set measurement parameters (Figure 2H): Go to Analyze > Set Measurements…
  a. In the pop-up window, select Area, Mean gray value, Centroid, Stack position, and Display label.
  b. Deselect all other parameters.
  c. Set Redirect to None and adjust the decimal places as preferred.
  d. Click OK.
48. Create a new macro in Fiji/ImageJ by going to File > New > Script. **Note:** If the macro is already saved, open it by dragging the macro file into the taskbar in Fiji/ImageJ, and skip to step 51.
49. Define the coding language as “ImageJ Macro” in the macro window by navigating to Language > ImageJ Macro.
50. Copy the following code into the script editor.

~~~
>//Set the segmentation directory as the location that the ROI zip file is
saved in
>#@ File (label = “Segmentation directory”, style = “directory”) output
>//Prompts the user to assign the number channel that each junction
protein is in
>#@ String (label = “ZO-1”, value = ““) ZO1
>#@ String (label = “E-cadherin”, value = ““) Ecad
>//Assigns the directory location of the image to the variable “dir”
>dir = getDir(“image”);
>//Assigns the filename of the image to the variable “iname”
>iname = getInfo(“image.filename”);
>selectWindow(iname);
>run(“ROI Manager…”);
>roiManager(“Open”, output + “/” + “ZO1_” + iname + “_BOND.zip”)
>//Renames the ROIs in ascending order. Ex: ROI1, ROI2, ROI3, etc.
>numROIs = roiManager(“count”);
>for (i = 0; i < numROIs; i++) {
      >roiManager(“Select”, i);
      >roiManager(“Rename”, “ROI_” + (i + 1)); }
>run(“Split Channels”);
>if (ZO1 > 0) {
      >selectImage(“C” + ZO1 + “-” + iname);
      >run(“Clear Results”);
      >// loop through ROIs
      >for(i = 0; i < numROIs; i++) {
             >roiManager(“Select”, i);
             >if (is(“line”)) {
                 >run(“Line to Area”); }
             >for (u = 1; u <= nSlices; u++) {
                 >setSlice(u);
                 >run(“Measure”); }}
      >saveAs(“Results”, dir + “_ZO1_” + iname + “_BOND.csv”); }
>if (Ecad > 0) {
   >selectImage(“C” + Ecad + “-” + iname);
   >run(“Clear Results”);
   >// loop through ROIs
   >for(i=0; i<numROIs;i++) {
      >roiManager(“Select”, i);
      >if (is(“line”)) {
        >run(“Line to Area”); }
      >for (u = 1; u <= nSlices; u++) {
         >setSlice(u);
  >run(“Measure”); }}
  >saveAs(“Results”, dir + “_Ecad_” + iname + “_BOND.csv”);
>}
~~~

51. Make any necessary adjustments (e.g. modify labels for ZO-1 if quantifying a different junctional protein), and save the new script for future use by navigating to File > Save As… in the open macro window
  a. Save the script to the quantification folder (Figure 2I, red box #1).
52. Execute the script by clicking the “Run” button at the bottom of the macro screen (Figure 2I, red box #2). **Note:** User comments provide detailed, line-by-line explanations of the code and are indicated by “//”. These comments serve as annotations to enhance readability and will not interfere with code execution. **Critical:** The segmentation directory must be set to the file location of the TIFF image being quantified.
  a. A prompt window will appear in Fiji/ImageJ, allowing the user to specify the segmentation directory, which serves as the destination folder for the quantification files. Additionally, the user will designate the specific channels in the original image to be quantified.
  b. The order of the channels can be found with the channel (C) slider in the original image (Figure 2G).
  c. The code will open the ROI zip file ONLY if it was saved with the required filename format of “ZO1_imagefilename.tif_BOND.zip” (Figure 2F).
  d. Each channel will be separated into individual images from the original image, and then the intensity of ZO-1 and E-cadherin across all Z slices within each ROI will be measured (Table 3 and Table 4).
  e. The results for ZO-1 and E-cadherin quantification will automatically be saved as individual CSV files to the quantification folder (Figure 2J).

**Table 3:** Example of raw ZO-1 intensity data from vertical junction quantification in Fiji/ImageJ. *Contents of column abbreviated to save space in data table. In raw data, it appears as: C2-CTRL.tif:c:2/4 z:1/30 - _CTRL_2_

|  | Label* | Area | Mean | X | Y | Slice |
| --- | --- | --- | --- | --- | --- | --- |
| 1 | CTRL | 1.513 | 0.516 | 3.986 | 25.383 | 1 |
| 2 | CTRL | 1.513 | 0.527 | 3.986 | 25.383 | 2 |
| 3 | CTRL | 1.513 | 0.704 | 3.986 | 25.383 | 3 |
| 4 | CTRL | 1.513 | 0.828 | 3.986 | 25.383 | 4 |
| 5 | CTRL | 1.513 | 1.054 | 3.986 | 25.383 | 5 |
| 6 | CTRL | 1.513 | 1.452 | 3.986 | 25.383 | 6 |
| 7 | CTRL | 1.513 | 2.656 | 3.986 | 25.383 | 7 |
| 8 | CTRL | 1.513 | 5.371 | 3.986 | 25.383 | 8 |
| 9 | CTRL | 1.513 | 15.161 | 3.986 | 25.383 | 9 |
| 10 | CTRL | 1.513 | 37.742 | 3.986 | 25.383 | 10 |
| 11 | CTRL | 1.513 | 54.199 | 3.986 | 25.383 | 11 |
| 12 | CTRL | 1.513 | 33.93 | 3.986 | 25.383 | 12 |
| 13 | CTRL | 1.513 | 13.392 | 3.986 | 25.383 | 13 |
| 14 | CTRL | 1.513 | 8.71 | 3.986 | 25.383 | 14 |
| 15 | CTRL | 1.513 | 6.677 | 3.986 | 25.383 | 15 |
| 16 | CTRL | 1.513 | 5.177 | 3.986 | 25.383 | 16 |
| 17 | CTRL | 1.513 | 4.866 | 3.986 | 25.383 | 17 |
| 18 | CTRL | 1.513 | 4.253 | 3.986 | 25.383 | 18 |
| 19 | CTRL | 1.513 | 3.699 | 3.986 | 25.383 | 19 |
| 20 | CTRL | 1.537 | 0.582 | 6.366 | 36.74 | 1 |
| 21 | CTRL | 1.537 | 0.767 | 6.366 | 36.74 | 2 |
| 22 | CTRL | 1.537 | 0.878 | 6.366 | 36.74 | 3 |
| 23 | CTRL | 1.537 | 1.095 | 6.366 | 36.74 | 4 |
| 24 | CTRL | 1.537 | 1.529 | 6.366 | 36.74 | 5 |
| 25 | CTRL | 1.537 | 2.884 | 6.366 | 36.74 | 6 |
| 26 | CTRL | 1.537 | 6.989 | 6.366 | 36.74 | 7 |
| 27 | CTRL | 1.537 | 19.265 | 6.366 | 36.74 | 8 |
| 28 | CTRL | 1.537 | 45.429 | 6.366 | 36.74 | 9 |
| 29 | CTRL | 1.537 | 50.857 | 6.366 | 36.74 | 10 |
| 30 | CTRL | 1.537 | 37.36 | 6.366 | 36.74 | 11 |
| 31 | CTRL | 1.537 | 20.497 | 6.366 | 36.74 | 12 |
| 32 | CTRL | 1.537 | 9.534 | 6.366 | 36.74 | 13 |
| 33 | CTRL | 1.537 | 6.921 | 6.366 | 36.74 | 14 |
| 34 | CTRL | 1.537 | 5.312 | 6.366 | 36.74 | 15 |
| 35 | CTRL | 1.537 | 4.852 | 6.366 | 36.74 | 16 |
| 36 | CTRL | 1.537 | 4.339 | 6.366 | 36.74 | 17 |
| 37 | CTRL | 1.537 | 4.032 | 6.366 | 36.74 | 18 |
| 38 | CTRL | 1.537 | 3.492 | 6.366 | 36.74 | 19 |

**Table 4:** Example of raw E-cadherin intensity data from vertical junction quantification in Fiji/ImageJ. *Contents of column abbreviated to save space in data table. In raw data, it appears as: C2-CTRL.tif:c:2/4 z:1/30 - _CTRL_2_

|  | Label* | Area | Mean | X | Y | Slice |
| --- | --- | --- | --- | --- | --- | --- |
| 1 | CTRL | 1.513 | 0.758 | 3.986 | 25.383 | 1 |
| 2 | CTRL | 1.513 | 0.769 | 3.986 | 25.383 | 2 |
| 3 | CTRL | 1.513 | 0.844 | 3.986 | 25.383 | 3 |
| 4 | CTRL | 1.513 | 0.892 | 3.986 | 25.383 | 4 |
| 5 | CTRL | 1.513 | 1.204 | 3.986 | 25.383 | 5 |
| 6 | CTRL | 1.513 | 1.742 | 3.986 | 25.383 | 6 |
| 7 | CTRL | 1.513 | 2.199 | 3.986 | 25.383 | 7 |
| 8 | CTRL | 1.513 | 2.876 | 3.986 | 25.383 | 8 |
| 9 | CTRL | 1.513 | 4.419 | 3.986 | 25.383 | 9 |
| 10 | CTRL | 1.513 | 5.801 | 3.986 | 25.383 | 10 |
| 11 | CTRL | 1.513 | 8.188 | 3.986 | 25.383 | 11 |
| 12 | CTRL | 1.513 | 10.634 | 3.986 | 25.383 | 12 |
| 13 | CTRL | 1.513 | 11.511 | 3.986 | 25.383 | 13 |
| 14 | CTRL | 1.513 | 11.016 | 3.986 | 25.383 | 14 |
| 15 | CTRL | 1.513 | 9.952 | 3.986 | 25.383 | 15 |
| 16 | CTRL | 1.513 | 8.398 | 3.986 | 25.383 | 16 |
| 17 | CTRL | 1.513 | 7.833 | 3.986 | 25.383 | 17 |
| 18 | CTRL | 1.513 | 7.07 | 3.986 | 25.383 | 18 |
| 19 | CTRL | 1.513 | 6.285 | 3.986 | 25.383 | 19 |
| 20 | CTRL | 1.537 | 0.757 | 6.366 | 36.74 | 1 |
| 21 | CTRL | 1.537 | 0.884 | 6.366 | 36.74 | 2 |
| 22 | CTRL | 1.537 | 1.138 | 6.366 | 36.74 | 3 |
| 23 | CTRL | 1.537 | 1.386 | 6.366 | 36.74 | 4 |
| 24 | CTRL | 1.537 | 1.815 | 6.366 | 36.74 | 5 |
| 25 | CTRL | 1.537 | 2.376 | 6.366 | 36.74 | 6 |
| 26 | CTRL | 1.537 | 3.302 | 6.366 | 36.74 | 7 |
| 27 | CTRL | 1.537 | 4.794 | 6.366 | 36.74 | 8 |
| 28 | CTRL | 1.537 | 6.968 | 6.366 | 36.74 | 9 |
| 29 | CTRL | 1.537 | 9.079 | 6.366 | 36.74 | 10 |
| 30 | CTRL | 1.537 | 10.566 | 6.366 | 36.74 | 11 |
| 31 | CTRL | 1.537 | 11.111 | 6.366 | 36.74 | 12 |
| 32 | CTRL | 1.537 | 11.095 | 6.366 | 36.74 | 13 |
| 33 | CTRL | 1.537 | 12.249 | 6.366 | 36.74 | 14 |
| 34 | CTRL | 1.537 | 11.825 | 6.366 | 36.74 | 15 |
| 35 | CTRL | 1.537 | 11.714 | 6.366 | 36.74 | 16 |
| 36 | CTRL | 1.537 | 11.63 | 6.366 | 36.74 | 17 |
| 37 | CTRL | 1.537 | 11.005 | 6.366 | 36.74 | 18 |
| 38 | CTRL | 1.537 | 10.476 | 6.366 | 36.74 | 19 |

**Table 5:**
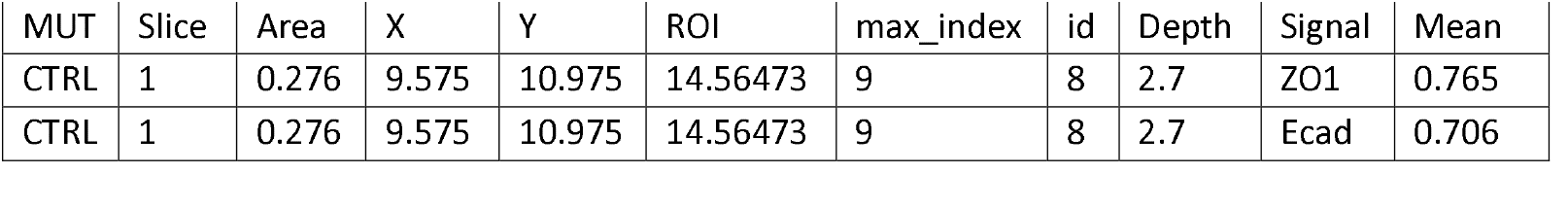
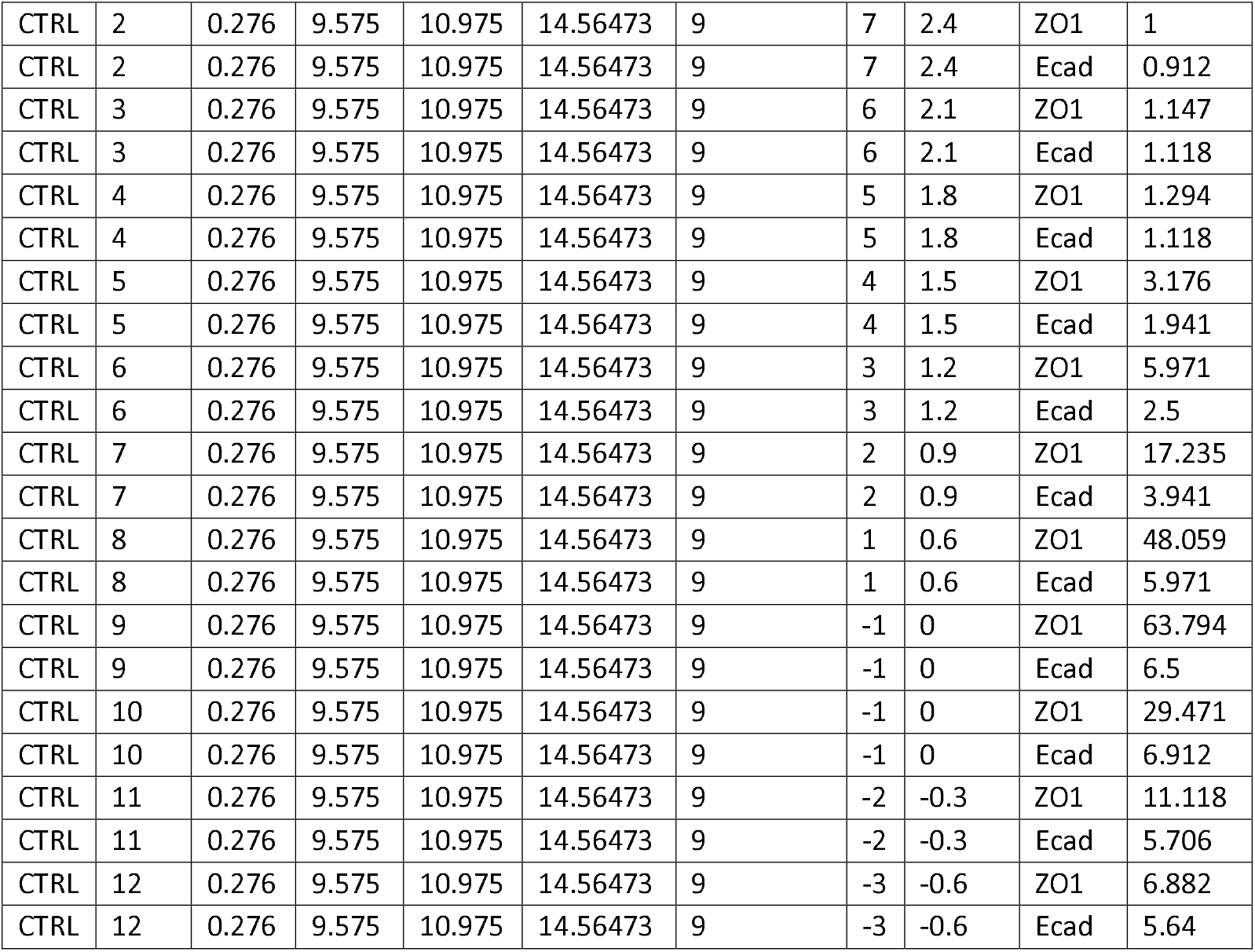
Example of “filtered” dataframe output from RStudio compilation and analysis of vertical junction profiling.

**Note:** Sample datasets generated using the Fiji/ImageJ vertical junction quantification protocol are provided in Tables 3 and 4. As described in step-by-step method details, step 52, the code outputs individual files for each junctional protein. Each segmented junction is represented by multiple rows of data, corresponding to individual Z-slices, which results in large datasets. To show the output format in a manageable space, we included data from 19 Z-slices across two segmented junctions as an example.

### Vertical profiling: Data analysis and graphing with R Studio

#### Timing: 10 minutes

Here, we describe a method utilizing the R programming language to compile junction intensity data previously obtained from the Fiji/ImageJ portion of the protocol (Steps 26-52). This approach generates a graph that visualizes and compares the intensity profiles of each measured junction marker across the z-stack.

The layout of RStudio differs from that of the Fiji/ImageJ macro window. In RStudio, the Source pane (typically in the upper left) is where code is written, edited, and executed. The Console, located in the bottom left, displays real-time execution of commands and maintains a history of previously run code within the session. Any errors that occur will be shown in this window. A “>” prompt in the Console indicates that the code has executed successfully, and that R is ready for the next command. The Environment pane, found in the upper right, contains active dataframes created during the session, which can be viewed at any time by clicking on them. Lastly, the Plots pane in the bottom right displays graphs generated by the code.

53. Download and install the R programming language along with the RStudio software interface.
54. Open RStudio and create a new script by navigating to File > New File > R Script. Save the script in the same directory as the ZO-1 and E-cadherin CSV files. **Note:** To run the RStudio code line by line, use the “Ctrl + Enter” keys. User comments provide detailed, line-by-line explanations of the code and are indicated by “#”. These comments serve as annotations to enhance readability and will not interfere with code execution.
55. Use the following code to load the necessary R packages:

~~~
>pacman::p_load(ggsci,rstudioapi,tidyverse, wesanderson, Rmisc,
ggplot2, plotly, carData, car, inferr, agricolae, multcompView, dplyr,
plyr, multcomp, corrplot, gridExtra, grid, ggpubr, ClusterRankTest,
readr,viridis, ggforce, purrr, ggdark)
~~~

**Note:** The packages need to be loaded only once per RStudio session. Each package has specific functions and syntax for data analysis.

56. Set the working directory to location of the script using the following commands:

~~~
>setwd(dirname(rstudioapi::getActiveDocumentContext()$path))
>dir <-getwd()
~~~

57. Create a dataframe named CSV that contains the Slice, Area, X, Y, and mean intensity values for each ROI in the ZO-1 quantification file. **Note:** The “%>%” (pipe) operator, when placed at the end of a line of code, directs the operation to continue onto the next line while still working within the same dataframe. Omitting this operator requires the user to specify the dataframe (e.g., the CSV dataframe) at the beginning of each line of code. When the following code is executed, a message detailing the column specifications will be displayed in the console. This is a standard output and should not be mistaken for an error message.

~~~
>#Identify all ZO-1 csv files in the folder
>CSV <- list.files(path = dir, pattern = “_ZO1.+.csv”, full.names = TRUE, recursive = TRUE) %>%
 >#reading the path to your CSV files
 >map_dfr(read_csv, .id = “Path”)%>%
 >#separates these columns out of the raw data
 >dplyr::select(Label, Slice, Area, X, Y, Mean) %>%
 >#Changing the mean (average pixel intensity) column to be renamed as ZO1
 >dplyr::rename(ZO1 = Mean)
~~~

58. Construct a dataframe named “Ecad” to list the average pixel intensity for each slice and ROI from all E-cadherin .CSV files.

~~~
>Ecad <-list.files(path = dir, pattern = “_Ecad.+.csv”, full.names =
TRUE, recursive = TRUE) %>%
   >map_dfr(read_csv, .id = “Path”) %>%
   >dplyr::select(Mean)%>%
   >dplyr::rename(Ecad = Mean)
~~~

59. Merge the two data sets by adding the E-cadherin mean intensity column to the existing “CSV” dataframe, which contains ZO-1 measurements. **Critical:** The label separation portion of the code will need to be edited based on information found under the “Label” column within the “CSV” dataframe.
  a. The Label column within the dataset will be separated into distinct categories to organize by experiment, mutation, field of view (FOV), etc. The name is separated based on the placement of the underscores (“_”). Any entries labeled as “NA” will be excluded from the defined categories. For example, if the label name is “C2-CTRL.tif:c:2/4 z:1/30 - _CTRL_2_,” the following code will categorize “CTRL” as the experimental condition/mutant “MUT,” and “2” as “FOV,” while excluding the remainder of the label name.
  b. The code creates a unique identifier for each ROI segmented in Tissue Analyzer by using the X and Y coordinate values (sqrt(X^2+Y^2)). This value will be stored as a character string, enabling it to function as a label rather than a numeric value.

~~~
>data<-CSV %>% bind_cols(Ecad)%>%
>#separates Label column into different components based on “_”
 >separate(Label, c(NA, “MUT”, “FOV”, NA), sep=“_”) % >%
 >#Creates an “ROI value” based on the X and Y measurements
 >mutate( ROI = sqrt(X^2+Y^2))
 >#stores the new ROI values as character strings
 >data$ROI <-as.character(data$ROI)
~~~

60. Run the following code to align the slices containing the brightest ZO-1 intensity for each ROI, and compile intensity data for regions located up to 1.5 µm apical and 3 µm basal relative to the brightest ZO-1 slice. **Note:** If the z-stack is acquired from the top to the bottom of the cell, lower slice numbers (e.g., slice 2) correspond to apical regions, while higher slice numbers (e.g., slice 30) correspond to basal regions.
  a. The user defines a slicing range based on cell height and the distribution of the junctional proteins of interest. For example, with a pixel depth of 0.3 µm during MDCK image acquisition, the normalized slicing range is defined as 5 slices (1.5 µm) apical and 10 slices (3 µm) basal of the brightest ZO-1 slice for each ROI.
  b. The original slice number containing the brightest ZO-1 intensity for each ROI is aligned by assigning an ID based on the row number in the newly sliced stack.
  c. The ID is converted to microns using the pixel depth of the image, which may vary depending on acquisition parameters. **Critical:** The pixel depth of the image can be determined in Fiji/ImageJ. Open the image, then navigate to Image > Properties. The number listed next to “Voxel depth” is the number that should be used in the depth conversion step of the code.
  d. To simplify graphing in R, a new column is created to label the signal type (e.g., “Ecad” or “ZO1”), and the corresponding intensity values are displayed in a separate column for each slice.

~~~
>#Groups the data by specified categories: MUT, FOV, and ROI
>filtered <-data %>% group_by(MUT, FOV, ROI) %>%
 >#Identifies the row with the maximum ZO-1 intensity for each ROI
 >mutate(maxZO1 = which.max(ZO1),
 #Calculates the number of slices after the brightest ZO-1 slice
 lastZO1 = n() - maxZO1) %>%
 >#Excludes ROIs where the brightest ZO-1 measurement is below slice 5
 >filter(maxZO1 >= 5) %>%
 >#Excludes ROIs with less than 10 slices basal from the brightest ZO-1
 >filter(lastZO1 >= 10) %>%
 >#Retains slices within 5 apical and 10 basal slices of the brightest
ZO-1 intensity for each ROI
 >slice((which.max(ZO1) - 5):(which.max(ZO1) + 10)) %>%
 >#Adds a new column “id” stating the row position within the new stack
 >mutate(id = -(row_number()),
 #Converts pixel depth to microns (adjust scaling factor as needed)
 Depth = ((id * 0.3) + 0.3)) %>%
 >#Reshapes the data to include columns for signal type and
corresponding intensity values
 >pivot_longer(ZO1:Ecad, names_to = “Signal”, values_to = “Mean”)
~~~

61. Execute the following code to calculate the average and minimum intensities for each signal type across the normalized stack.

~~~
>#Calculates the average intensity and minimum intensity of each signal
type at each depth for each condition
>CSVMean <-filtered %>% group_by(Signal, MUT, Depth)%>%
 >summarise(Mean=mean(Mean),
        Min=min(Mean))
>#Extracts the maximum and minimum Ecad intensity values for the CTRL condition from the previously calculated averages
>CSVMaxEcad <-CSVMean %>% group_by(Signal, MUT)%>%
 >summarise(Max=max(Mean),
       Min=min(Min)) %>%
 >filter(MUT==“CTRL” & Signal==“Ecad”)
>CSVMaxZO1 <-CSVMean %>% group_by(Signal, MUT)%>%
 >summarise(Max=max(Mean),
       Min=min(Min)) %>%
>filter(MUT==“CTRL” & Signal==“ZO1”)
~~~

62. Run the following code to perform min-max normalization on the CTRL condition, which scales the maximum signal intensity to 1 and the minimum to 0. This normalization allows direct comparison of signal intensities across different experimental conditions and signal types when graphed.

~~~
#Normalize the maximum intensity for each signal in the CTRL condition to
be equal to one
>filtered <-filtered %>% group_by(Signal)%>%
   >mutate( Mean = case_when( Signal==“ZO1” ~ (Mean - CSVMaxZO1$Min)/(
CSVMaxZO1$Max-CSVMaxZO1$Min),
               Signal==“Ecad” ~ (Mean - CSVMaxEcad$Min)/(
CSVMaxEcad$Max-CSVMaxEcad$Min)))
>#Places the different conditions in the order to be plotted later
>filtered$MUT <-factor(filtered$MUT, levels = c(“CTRL”, “KD”, “Condition
3”))
>#Organizes ZO1 to be before Ecad when graphing
>filtered$Signal <-factor(filtered$Signal, levels = c(“ZO1”, “Ecad”))
~~~

63. Use the following code to configure a graph with an x-axis ranging from −0.1 to 1.5 and a y-axis scaled according to Depth values. A vertical grey line will be included for reference at x = 1.

~~~
>#Define general aesthetic for the graph
>plot <-ggplot(filtered, aes(x=Mean, y=Depth)) +
  facet_grid(rows=vars(MUT))+
  geom_point(aes(color=Signal), alpha=0)+
  theme_classic()+
  geom_vline(xintercept = 1, colour = “grey70”, size=1.25)+
  scale_shape_manual(values=c(3,4,8,15,16,17,18,19))+
  scale_color_manual(values=c(“magenta”,”green”))+
  scale_fill_manual(values=c(“magenta”,”green”))+
  xlim(−0.1,1.5)+
  xlab(“Relative intensity (a.u.)”)+
  ylab(“Depth from apical (Âµm)”)+
  guides(shape=FALSE)+
  guides(color=FALSE)+
  guides(fill=FALSE)
>#Plot the base for the graph
>plot
~~~

64. Generate a plot to visualize the intensity distribution and standard deviation for each signal type on the base graph.

~~~
>#Calculates standard error for the defined categories
>Meansum <-filtered %>% group_by(MUT,Signal,Depth) %>%
  >dplyr::summarise(sd=sd(Mean),
   >n = n(), #Calculate sample size
   >SE = sd /sqrt(n), #Compute standard Error
         Mean=mean(Mean))
>#Adds the distribution of the average signal values to the base graph
>plot2 <-plot +
 geom_path(data = Meansum, aes(color=Signal), size=1)
>#Adds the standard deviation as a dashed line around the signal values
>plot2 +
 geom_ribbon(data = Meansum, aes(fill=Signal, xmax = Mean+SE, xmin = Mean-
SE), alpha=0.2) +
 geom_path(data = Meansum, aes(y=Depth, x=Mean+SE, color=Signal), linetype=“dashed”) +
 geom_path(data = Meansum, aes(y=Depth, x=Mean-SE, color=Signal), linetype=“dashed”)
~~~

65. In the Plots pane, navigate to Export > Save as Image… and save the graph to the desired location.

### Expected outcomes

The horizontal junction analysis of the provided test files (Supp. file 1-2) in Fiji/ImageJ will generate datasets that are similar as the examples shown in Table 1 and Table 2 (one individual .CSV file per ROI). This allows for a quantitatively comparison between the fluorescence intensity of junctional markers and a protein of interest. The following processing in R Studio will generate standardized and normalized fluorescence profiles for comparison across conditions and graphical visualization of the dataset, as illustrated in Figure 3.

**Figure 3:**
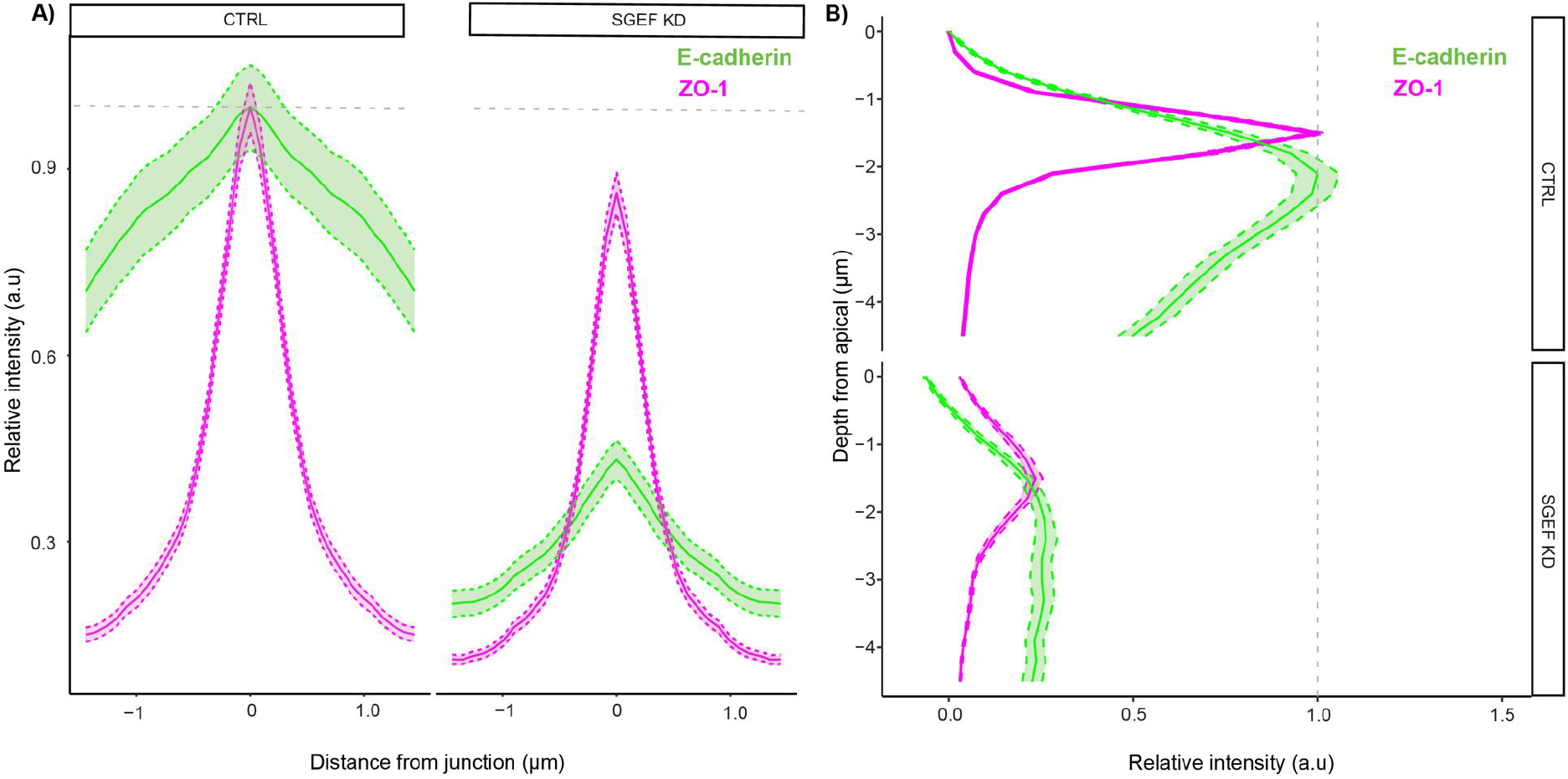
Individual plots of horizontal and vertical junction analysis data (A-B) Plots of horizontal junction data acquired from step-by-step method details, step 24, comparing CTRL and KD conditions. (C) Plot of vertical junction data acquired from step-by-step method details, step 63, comparing CTRL and KD conditions. The figure contains data from one field of view per condition, and therefore, there is a large amount of variability in the spread of numbers. The magenta line represents the average vertical distribution of ZO-1 intensity across all the junctions, while the green line represents that of E-cadherin intensity.

In the vertical junction analysis, after processing individual junction ROIs using Tissue Analyzer in Fiji/ImageJ, the resulting dataset should resemble the examples shown in Table 3 and Table 4 for ZO-1 and E-cadherin values respectively. A distinctive feature of this protocol is the ability to generate a graph that displays the intensity distributions of multiple junctional markers within the same plot. These graphs allow for visualization of the average intensity at individual junctions along with the variability within the dataset, as illustrated in Figure 3.

Comparison of these graphs across experimental conditions shows differences in the localization and intensity of E-cadherin and ZO-1. Specifically, KD cells exhibit reduced levels of both proteins, with a more diffuse distribution throughout the cell, suggesting a potential impairment of cell-cell junction integrity. Given that the expression levels of some junctional proteins can vary across experiments, we recommend confirming protein levels via Western blot before drawing definitive conclusions from intensity-based quantification.

### Quantification and statistical analysis

The quantification procedures are described in detail in the step-by-step method details, step 1-65. Statistical analyses can be done using R Studio, GraphPad Prism, or other appropriate statistical software. In Rabino et al. (2024), we first tested the data for normality and homogeneity of variance using the Shapiro-Wilk and Levene’s tests, respectively. When both assumptions were met, group means were compared using a one-way ANOVA followed by Tukey’s post-hoc test. If either assumption was not passed, a Kruskal-wallis test with a Wilcoxon post-hoc analysis was applied. Sample images are provided for running a complete test of both image analysis protocols (Supp. file 1-2).

### Limitations

There are a few limitations that should be considered when doing the horizontal junction quantification. Manual ROI selection may introduce user bias, affecting reproducibility, so it is important to be consistent when positioning ROIs in different junctions (e.g. at the center point between two tricellular junctions). The length of the ROI is also an important factor, an ROI that results in less than 16 pixels long to each side of the brightest point will be discarded, so make sure that you draw an ROI long enough to avoid this.

Another challenge would be that if the basal part of the monolayer is more spread out than the apical side, that is usually more constricted, the junctions will be more diagonal. These junctions will introduce artifacts in fluorescence intensity measurements when making a Z-projection. This can be corrected by aligning the Z-plane based on the maximum fluorescence intensity.

The variable vertical distribution of ZO-1 across monolayers plated on coverslips creates a challenge in capturing the signal throughout the entire z plane during the slicing step in step-by-step methods details, step 60. To include the entire vertical signal, a substantial buffer space must be acquired both apically and basally to the monolayer when imaging. Additionally, the protocol relies on highly specific file names, which may introduce errors when quantifying multiple experiments. Lastly, while these methods can be adapted for batch processing, the requirement for manual adjustments in Tissue Analyzer limits fully automated high-throughput quantification.

### Troubleshooting

#### Problem 1

The horizontal analysis generated less measurements than the number of ROIs defined. This could happen if the line drawn is shorter than 32 pixels

#### Potential solution

When the code “filter” is run, ImageJ is going to filter all the results and keep the ROIs that have 16 pixels to each side of the brightest point of ZO-1 (or the marker used for reference), and eliminates all ROIs that don’t meet this requirement. To minimize this, make sure that all the ROIs are at least 10 µm long.

#### Problem 2

Junctions have gaps or are non-continuous in the max projection created in vertical junction analysis step-by-step method details, step 28.

#### Potential solution

This issue may arise if the entire monolayer is not captured during imaging. To ensure all junctions are included, it is recommended to acquire a few z-planes above and below the monolayer. Biologically, an immature monolayer could also contribute to this problem. In our experiments, MDCK cells are typically allowed to polarize for approximately four days, though this duration will vary between different cell lines. Additionally, different experimental conditions can disrupt junction integrity.

#### Problem 3

Image does not show up in Tissue Analyzer (step-by-step method details, step 33). This problem mainly arises from file directory limitations. Tissue Analyzer is unable to access university networks or excessively long file paths.

### Potential solution

To resolve this, all images and quantification data should be saved in an easily accessible location on the user’s hard drive, while avoiding long file names and paths. Once quantification is complete, the files can be relocated as needed.

#### Problem 4

Automated segmentation fails to recognize the majority of the junctions regardless of strong blur modification in Tissue Analyzer (step-by-step method details, step 34). The most likely cause of this issue is insufficient intensity or poor localization of the junctional protein signal.

### Potential solution

To improve segmentation, the image can be cropped into smaller quadrants using the rectangle tool in Fiji/ImageJ before inputting it into Tissue Analyzer, which may enhance accuracy. If this does not resolve the problem, the user may manually draw the segmented lines following the step-by-step method details, step 28. Alternatively, the experiment can be repeated using a stronger junctional protein marker or by optimizing staining and/or imaging conditions, such as antibody concentration, laser power, exposure time, and/or resolution.

#### Problem 5

The Fiji/ImageJ or RStudio code runs into unexpected errors.

#### Potential solution

The most common sources of unexpected errors when running code include extra or missing brackets, semicolons, pipes (%>%), spaces, or other symbols. When modifying the code to accommodate different experimental conditions, it is essential to maintain the exact placement and formatting of all symbols (especially the quotation marks) as specified in the protocol to ensure proper functionality.

#### Problem 6

In step-by-step method details, step 59, an error similar to the following appears in the RStudio Console:

~~~
  >Warning message:
  >Expected 4 pieces. Missing pieces filled with ‘NA’ in 2542 rows [1441,
  1442, 1443, 1444, 1445, 1446, 1447, 1448, 1449, 1450, 1451, 1452,
1453, 1454, 1455, 1456, 1457, 1458, 1459, 1460, …].
~~~

### Potential solution

This warning occurs when there is a conflict between the specified parameters for separating the “Label” name into different categories and the actual formatting of the “Label” name. To troubleshoot this issue, users should first examine the “CSV” dataframe, specifically the formatting of the contents of the “Label” column, to ensure that the number of underscores is exactly one less than the number of desired segments.

For example, in structuring the “separate” command based on CTRL labels, a typical label can follow the format: `“C2-CTRL.tif:c:2/4 z:2/30-_CTRL_2_.”` Here, the underscores define four pieces. However, when KD labels are introduced, such as `“C2-KD.tif:c:2/4 z:24/41 - KD_1_,”` a missing underscore disrupts the separation process. This issue can be resolved by either renaming the original acquired images and re-running the quantification, or by simply adjusting the label formatting within RStudio. The following bolded line of code, added in the designated location, inserts an underscore before “KD” in the label name, ensuring consistency across datasets:

~~~
>data<-CSV %>% bind_cols(Ecad)%>%
      >#separates Label column into different components based on “_”
      >mutate(Label = gsub(“(KD)”, “_\\1“, Label)) %>%
      >separate(Label, c(NA, “MUT”, “FOV”, NA), sep=“_”) %>%
~~~

#### Problem 7

In step-by-step method details, step 60, an error similar to the following appears in the RStudio Console:

~~~
>Error in slice():
>! Can’t compute indices.
>i In group 7: Label = “C2-CTRL.tif:c:2/4 z:15/30 - _CTRL_2_”.
>Caused by error in dots[[group]]:
>! subscript out of bounds
~~~

### Potential solution

This error occurs when the slicing range specified in the which.max(ZO1) line of code exceeds the available rows in the z-stack. To troubleshoot, first remove the pipe (%>%) from the following line of code:

~~~
>lastZO1 = n() - maxZO1) %>%
~~~

Next, run the code up to this point and inspect the “filtered” dataframe in the R Environment. Check the values assigned to maxZO1 and lastZO1, ensuring that maxZO1 is at least 5 and the value for lastZO1 is at least 10 (these are predefined values in this protocol, but you must verify based on your specific dataset). This will ensure that there are at least 5 rows apical and 10 rows basal of the brightest points in the “data” dataframe.

This error arises because R cannot slice rows that do not exist. Unless the brightest ZO-1 values for all ROIs are located in the same z-plane, do not define the slicing range as the entire stack, because some ROIs will inevitably fall outside this range. Once the issue is identified, reintroduce the pipe to restore proper code functionality.

## Resource availability

### Lead contact

Further information and requests for resources and reagents should be directed to and will be fulfilled by the lead contact, Dr. Rafael Garcia-Mata.

### Technical contact

Technical questions on executing this protocol should be directed to and will be answered by the technical contact, Madeline Lovejoy.

## Materials availability

This study did not generate new unique reagents.

## Data and code availability

The published article includes all datasets and code generated or analyzed during this study.

## Acknowledgments

This work was supported by the following grants from the National Institute of Health (NIH) awarded to R. Garcia-Mata (R01GM136826 and R15GM155874). The authors thank Lucia Gonzalez-Blotta from the University of Toledo for her contributions in testing the protocol and identifying potential challenges that may arise for other researchers.

## Author contributions

Conceptualization, A.R. and R.G.-M. Investigation, A.R. Methodology, A.R., M.L., and C.A.M. Formal analysis, A.R. and R.G.-M. Writing, M.L., C.A.M., and R.G.-M. Visualization, M.L. and C.A.M. Resources, R.G.-M. Funding acquisition, R.G.-M. Supervision, R.G.-M. Project administration, R.G.-M.

## Declaration of interests

The authors declare no competing interests.

## Notes

### Competing Interest Statement

The authors have declared no competing interest.

## References

Rabino, A., Awadia, S., Ali, N., Edson, A., and Garcia-Mata, R. (2024). The Scribble-SGEF-Dlg1 complex regulates E-cadherin and ZO-1 stability, turnover and transcription in epithelial cells. J Cell Sci 137. 10.1242/jcs.262181.

Aigouy, B., Umetsu, D., and Eaton, S. (2016). Segmentation and Quantitative Analysis of Epithelial Tissues. Methods Mol Biol 1478, 227–239. 10.1007/978-1-4939-6371-3_13.

Awadia, S., Huq, F., Arnold, T.R., Goicoechea, S.M., Sun, Y.J., Hou, T., Kreider-Letterman, G., Massimi, P., Banks, L., Fuentes, E.J., et al. (2019). SGEF forms a complex with Scribble and Dlg1 and regulates epithelial junctions and contractility. J Cell Biol 218, 2699–2725. 10.1083/jcb.201811114.

