## Supplementary figures and images for "Quantification of horizontal and vertical distribution of junctional proteins in fixed epithelial cells"

### Supp Fig 1

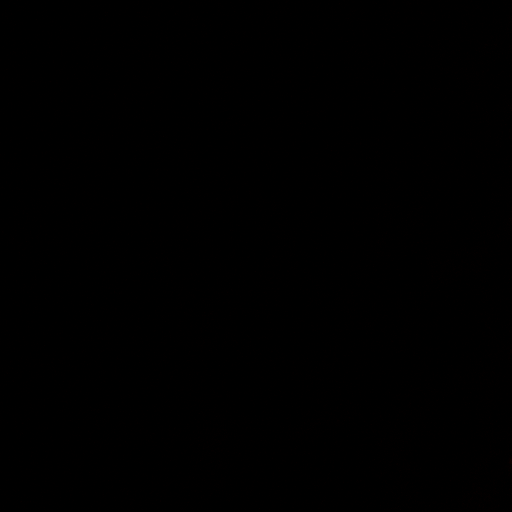

### Supp Fig 2

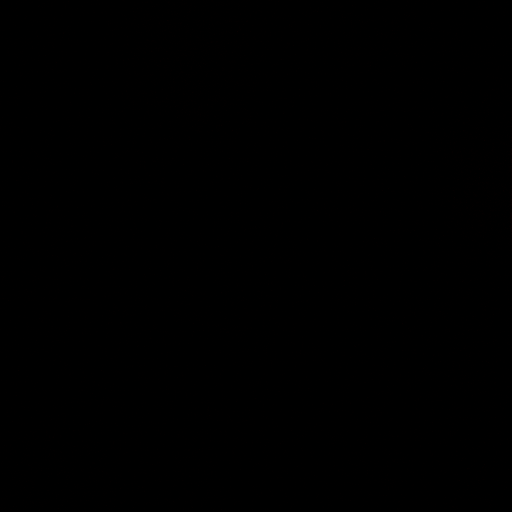
